# Mechanisms mediating arylidene-indolinones induced degradation: thoughts on “Discovery of a Drug-like, Natural Product-Inspired, DCAF11 Ligand Chemotype”

**DOI:** 10.1101/2024.03.05.582859

**Authors:** Chao Zhong, Ziying Wang, Zhaoyang Li, Haofeng Li, Qianhui Xu, Wanli Wu, Cong Liu, Yiyan Fei, Yu Ding, Boxun Lu

## Abstract

In the recent issue of Nature Communications (2023 Nov 30;14(1):7908), Xue et. al. reported a very interesting and significant discovery of a possible DCAF11 ligand chemotype that could be used as the “warhead” to design bifunctional compounds for targeted degradation via engaging the E3 ligase DCAF11 ^1^ (annotated as ref 1 hereafter). The discovery is of importance to the targeted protein degradation field and was inspired by previous reports suggesting that similar compounds may also engage the autophagosome protein LC3 for degradation and function as autophagy-tethering compounds (ATTECs) ^2, 3, 4, 5, 6, 7^, which seem to be inconsistent with ref 1. We think that the conclusions based on these data are not necessarily mutually exclusive. After performing additional experiments and analyses, we would like to discuss some possibilities explaining such discrepancies and make a few points of clarification.

## Results and analyses

Our points are described below based on analyses and new experimental results.

First, the chemical structures of all the compounds utilized to test LC3-binding are somewhat different from the original ones reported for degradation of mutant HTT proteins (mHTT) or for degradation of BRD4. In ref 1, compounds 5&18 were tested for LC3-binding (Supplementary Fig. 8e-f in ref 1). Noticeably, while these compounds have highly similar structures to the ones used in ATTEC studies, the substituent on the oxindole skeleton was switched from bromo to iodo in published ATTECs using GW5074 as the LC3B ligand ^2, 3, 4, 5, 6, 7^. In addition, based on the supplementary data in the publication, 1H Nuclear Magnetic Resonance (NMR) profiles of compounds 5&18 are different from the published ATTECs based on the supplementary information provided, likely because of differences in Z/E configuration.

Second, the LC3-binding assays utilized in ref 1 were different from the ones reported in our original study ^3^. Ref 1 states that there is no binding of the BODIPY-labelled compound 18 to LC3 in a fluorescence polarization (FP) experiment. In the presented data, the FP curve in ref 1 actually exhibited a change in polarization, albeit small, in response to different LC3 concentrations (cropped from Supplementary Fig. 8e in ref 1). The size of the response (~20 mP) is comparable to some of the other FP studies showing interactions ^8^. Typically, in fluorescence polarization (FP) experiments, the labeled ligand is maintained at a constant concentration below the dissociation constant (K_d_/10 or less), while the unlabeled molecule is titrated against it. The concentration range of the unlabeled molecule (LC3B protein in this case) should cover at least 100-fold below and 100-fold above the K_d 9_. This experimental condition allows the polarization signals to change from the lowest level when the labeled compound stays at an almost completely free state to the highest level when the labeled compound reaches a completely bound state, thereby maximizing the dynamic range of the experiment. In ref 1, a concentration of 100 μM was used for the labeled ligand (BODIPY-labelled compound 18). This concentration is likely much higher than the recommended concentration for FP experiments (K_d_/10 or less), potentially accounting for the small change of signals observed in the FP curve. Ref 1 also reported the conclusion that compound 5 does not bind to LC3B because it did not affect the thermal stability of purified recombinant LC3B (Supplementary Fig. 8f in ref 1). First, we’d like to reemphasize that compound 5 is not identical to GW5074 which we reported previously as an LC3B binder as we mentioned in the first point. In addition, it is also worth noting that in some cases, certain protein binders may only induce small or negligible thermal shifts ^10, 11^, making thermal detection alone insufficient to exclude the possibility of binding. Purified recombinant LC3B would also be devoid of any of the post-translational modifications that could affect the binding site and strength ^12^. Finally, the solubility of the compounds is poor, and this may further influence the compound-protein interaction measurements using these assays. Comparatively, the Oblique-Incidence Reflectivity Difference (OI-RD) assay utilized in the original ATTEC study measured the compound-protein interaction in a setting where the compounds were immobilized in a chip, minimizing the potential influence of poor solubility and the clustering tendency of the compounds ^3^. Consistent with this postulation, the surface immobilization of compounds with low solubility enables the detection of their protein-binding that is difficult to detect using other settings, as exemplified by the detection of the rapamycin-FRB interaction ^13^.

Third, to further clarify the issue with LC3B-binding, we performed additional Surface Plasmon Resonance (SPR), Bio-layer Interferometry (BLI), and 2D-NMR experiments, which all confirmed that the two original arylidene-indolinones reported as LC3B binders, GW5074 and AN1, do interact with LC3B (Fig. 1b-d). The affinities measured by SPR and BLI were weaker than the ones measured by OI-RD in the previous report, especially for GW5074, possibly because of the low solubility and clustering tendency of the compounds that may influence the affinity measurement by SPR and BLI. To further confirm the compound-LC3 binding in cells without the caveat of any missing binding-relevant post-translational modifications, we performed the cellular thermal shift (CETSA) assay, which confirmed a significant thermal shift of LC3B-I and LC3B-II in the cell lysates (Fig. 2). A recent preprint from Joshua A. Kritzer’s group also confirmed that GW5074 may interact with LC3B and GABARAP ^14^. We also agree with the conclusion that these LC3B-binders are probably not ideal candidates to inhibit autophagy, consistent with our previous results showing that they did not influence the global autophagy flux ^3^.

**Figure 1.**
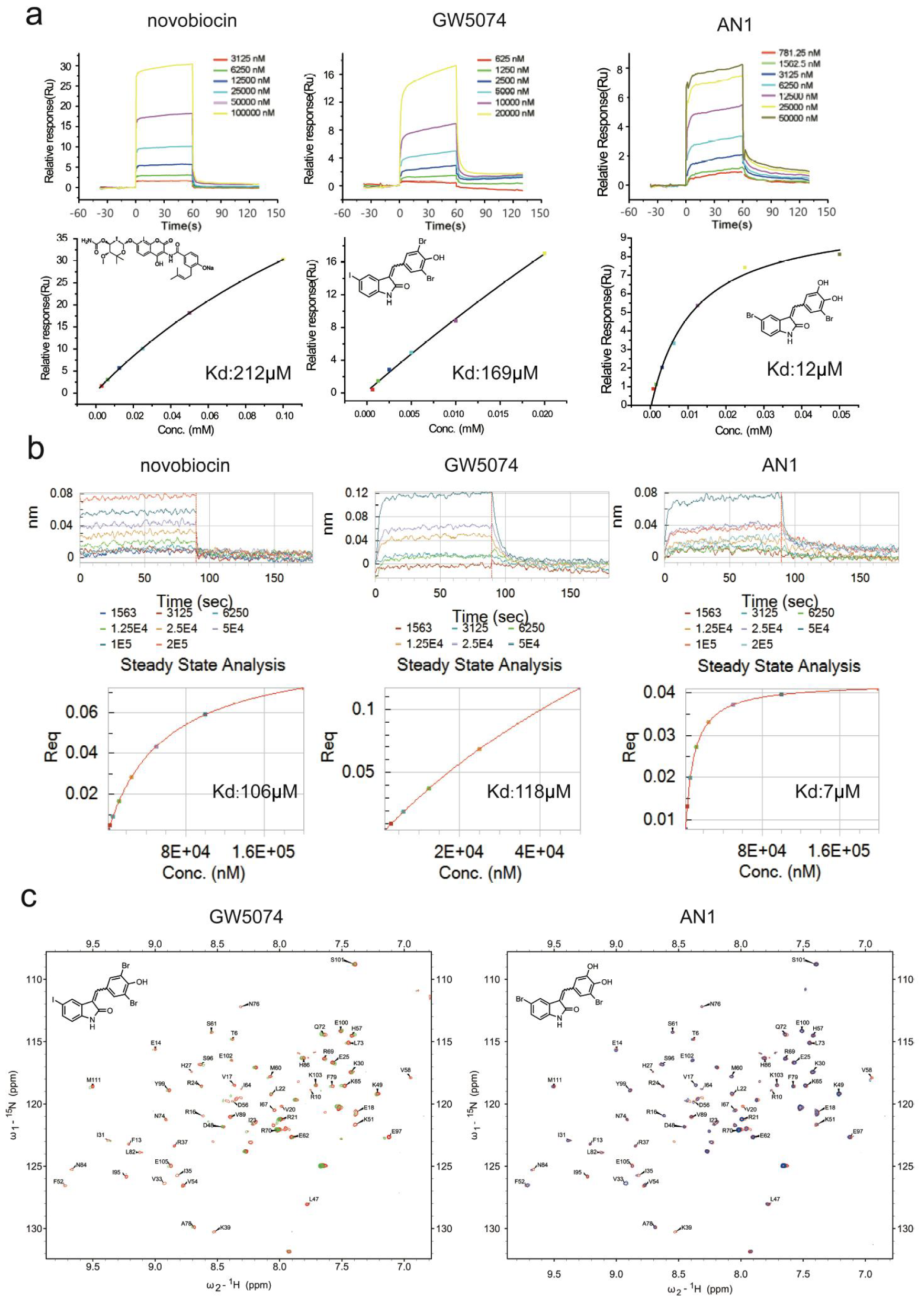
Confirmation of the interactions between LC3 and GW5074/AN1 *in vitro*. (a) Representative SPR assay results (up) and fitting curves with K_d_ values (down, affinity analysis of the steady-state response) for the indicated compound. LC3B was conjugated to the SPR chip for the measurement. Another previously reported LC3B binder with much higher solubility, novobiocin ^22^, was used as a positive control. (b) Similar to (a), but using BLI technology for the measurement. Req: response at equilibrium. (c) Representative (fingerprint) areas of LC3B 1H–15N HSQC spectra upon titration with GW5074 or AN1 (overlaid). Spectra of the free (red) and compound bound forms (green for GW5074 and orange for AN1) of LC3B are shown. Considering that novobiocin has a much better solubility (100 mg/mL) than GW5074, which also has a tendency to cluster in assay buffers, the LC3B-binding affinity of GW5074 is likely much higher than novobiocin and under-estimated by SPR (a) and BLI (b).

**Figure 2.**
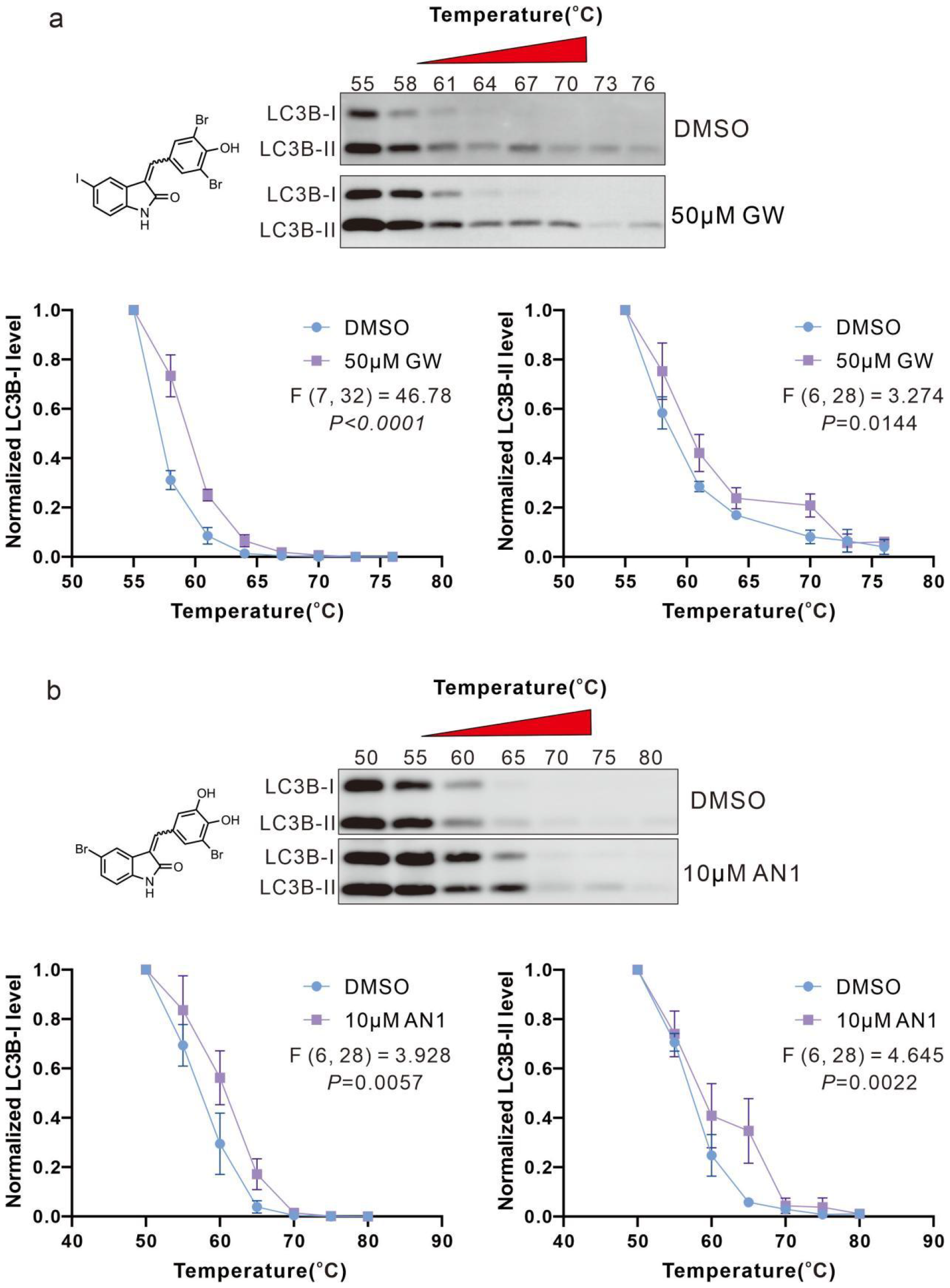
Cell Thermal Shift Assay (CETSA) assays validating the interactions between the indicated compounds and LC3B in cells. Representative Western blots and quantifications of the CETSA experiments using the 293T cell lysates incubated with GW5074 (GW, 50 μM) (a) or AN1(10 μM) (b) for 2 hours followed by heating for 5 minutes at the indicated temperatures. GW5074 and AN1 were shown to interact with both LC3B-I and LC3B-II based on this assay. The DMSO versus compound-treated samples were loaded in the same gels with the same exposure. We placed them up and down for easier comparison. 3 biological replicates were performed. Two-way ANOVA tests.

Fourth, while the detected LC3-binding affinities were not high in the SPR and BLI assays (~10 to ~100 μM Kd, Fig. 1), these compounds may still trigger the ternary complex formation and engage LC3B for the degradation, at least for certain specific targets. Firstly, the immunomodulatory drugs (IMiDs) that have been widely used in highly efficient (nM) PROTACs (PROteolysis TArgeting Chimeras) also exhibited >1 μ M Kd in the SPR assays ^15^, suggesting that affinities at this range may work for degraders. Secondly, the affinities of GW5074/AN1-LC3B interaction were possibly underestimated, as discussed earlier. Thirdly, while their DCAF11-binding affinities were not directly measured in ref 1, it seems that >10 μM compound concentrations were required to detect the interaction, which was still not saturated at 100 μM (Fig. 4f of ref 1), further supporting that the affinities in this range could be enough for degradation. Lastly and importantly, we directly confirmed that GW5074 and AN1 were able to induce the LC3B-mHTT interaction as detected by the OI-RD assay (Fig. 3).

**Figure 3.**
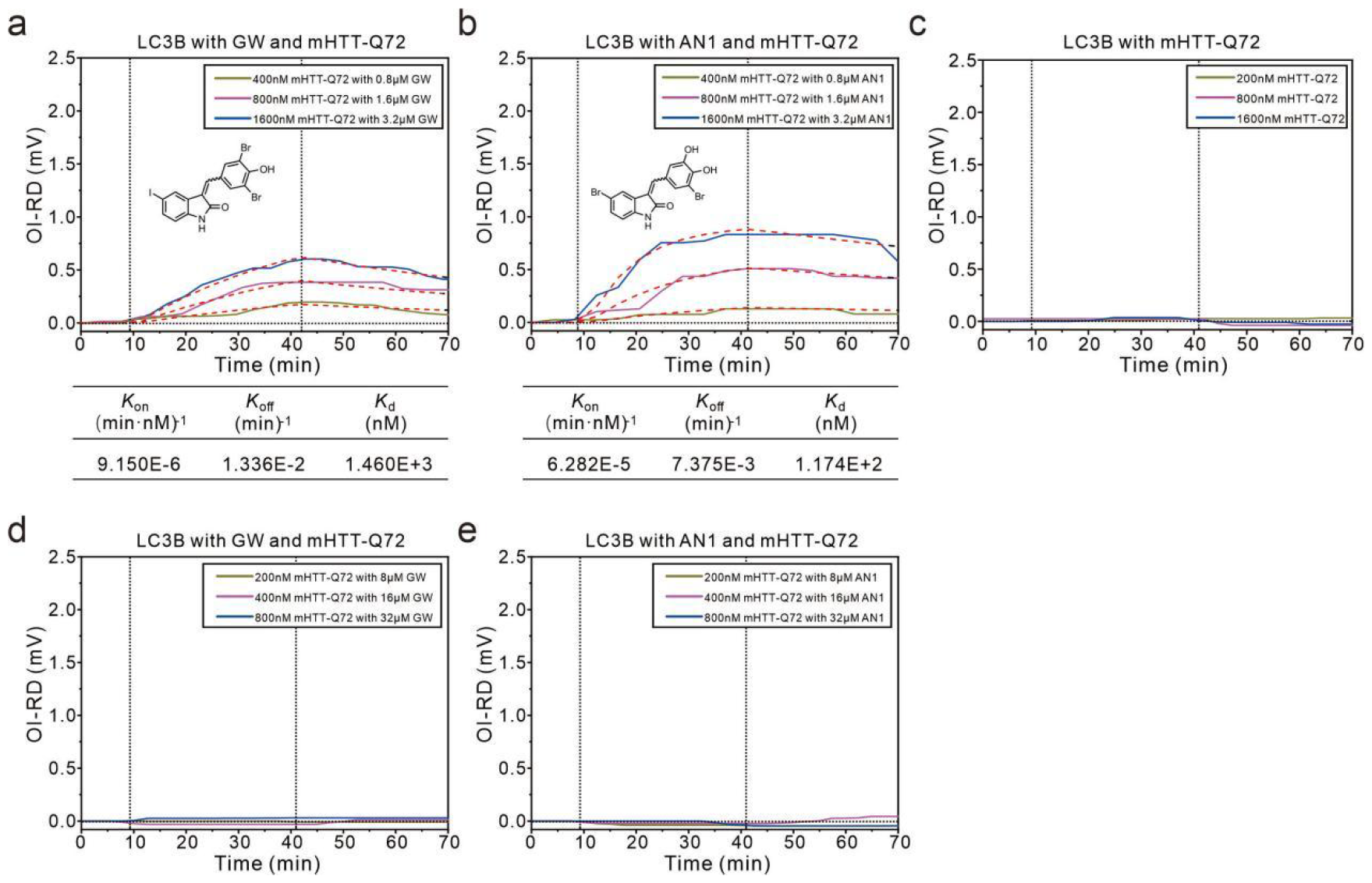
GW5074 and AN1 enhanced mHTT-LC3B interaction as detected by OI-RD. Association–dissociation curves of surface-immobilized recombinant purified LC3B protein with MBP-mHTTexon1-72Q (mHTT-Q72) with or without the indicated compounds at indicated concentrations. In association–dissociation curves, vertical dashed lines mark the start of the association and dissociation phases of the binding events. The red dashed curves are global fits by a Langmuir reaction model with the fitting parameters listed at the bottom of each plot. Note that the assay setting was different from the original ones we published previously in which the compounds rather than the proteins were immobilized. ~μM to sub-μM affinities between LC3B and mHTT-72Q were detected only when appropriate concentrations of GW5074 (a, GW) or AN1 (b) compound were present. No binding signals were observed for mHTT-72Q flowing through the LC3B-stamped chip alone in this assay with this setting (c). The detected binding signals were not from the compounds *per se* because their sizes are too small compared to the mHTT-72Q protein. When we increased the compound concentrations up to 40 times (d-e), no binding signals were detected, illustrating that the compounds themselves did not induce sufficient signals and the ternary complex formation could be disrupted by excessive compounds (hook effects).

**Figure 4.**
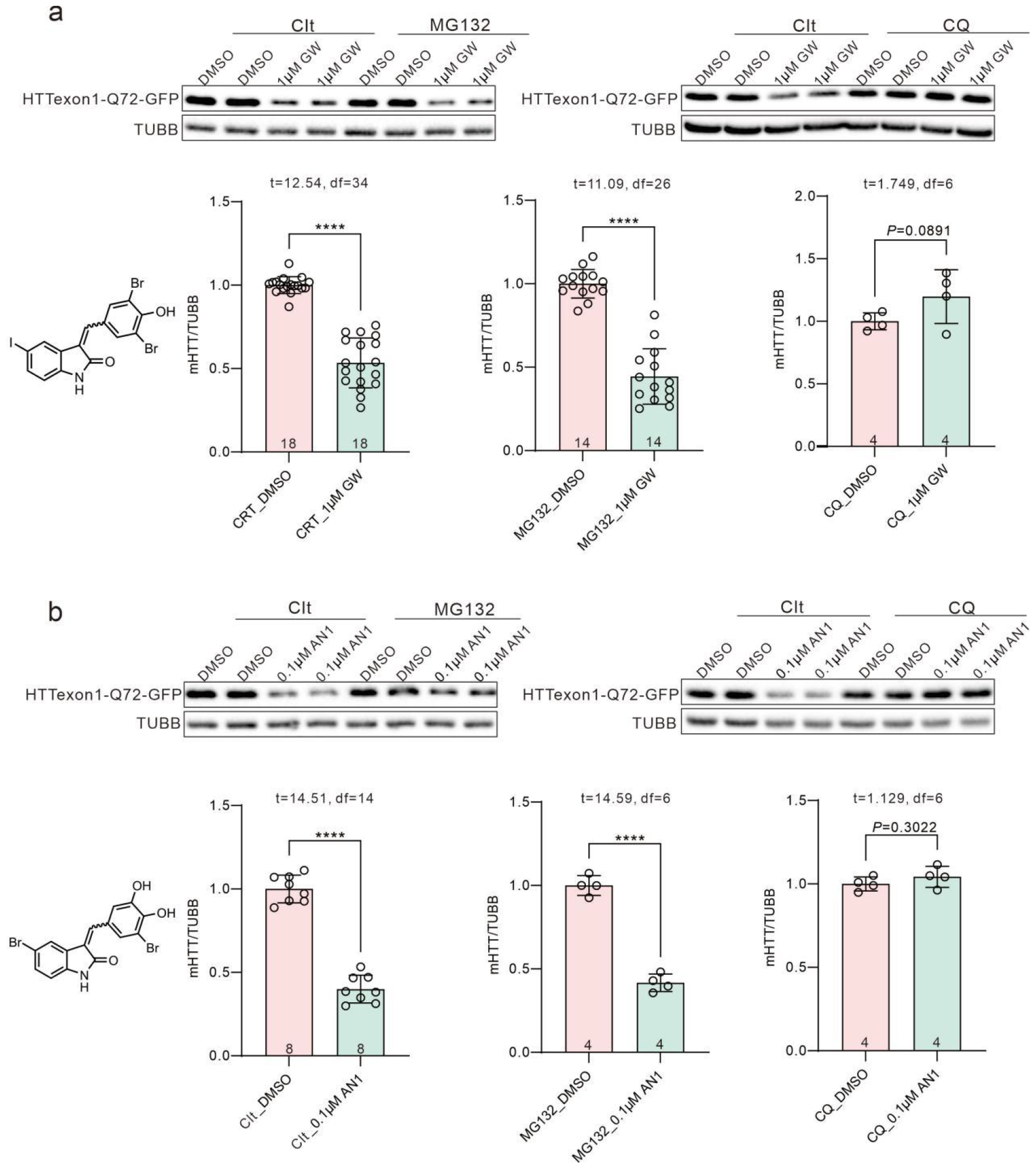
mHTT-lowering effects by GW5074 and AN1 were dependent on autophagy. Representative Western blots and quantifications of compound-treated HTTexon1-72Q-GFP overexpressed HEK293T cells. The cells were treated with the indicated compounds (1000 nM for GW5074 and 100 nM for AN1) with the indicated compounds (the proteasome inhibitor MG132, the autophagy inhibitor chloroquine (CQ), or equivalent DMSO control (Clt)). One-way ANOVA with post hoc Dunnett’s tests.

Fifth, we performed additional experiments to confirm that the degradation of mHTT with expanded polyQ by GW5074 or AN1 was mediated by autophagy. Since the primary cultured Huntington’s disease knock-in mouse neurons and patient fibroblasts utilized in the original ATTEC study are not easily accessible to many researchers, we tested the compounds’ effects on mHTT in 293T cells transiently transfected with the mHTT exon 1 fragment (Q72). Consistent with our previous studies, we observed the lowering of Q72 by the treatment of either GW5074 or AN1, and the degradation was inhibited by the autophagy inhibitor chloroquine (CQ) but not the proteasome inhibitor MG132 (Fig. 4). These experiments do not require any special materials and could easily be replicated. We did notice that the cell confluence may influence the compounds’ effects as typical for many cellular experiments and plating density at 8×10^5^/well in a 6-well plate gave the most robust degradation. The relatively high cell density may enhance the basal autophagy activity to degrade over-expressed mHTT.

Based on ref 1 and the analysis/discussion above, a likely scenario is that the arylidene-indolinones including GW5074, AN1, and their analogs, may interact with both LC3 and DCAF11. Their affinities to each protein could be influenced by subtle changes in the chemical group (such as iodo versus bromo), Z/E configuration, and/or the experimental conditions. The scenario becomes even more complicated in the cellular context, which may explain why these highly similar arylidene-indolinones may lead to different degradation mechanisms observed in different cell types on different targets. For example, for the same target PDEδ, the study in ref 1 and the ATTEC study utilized different cell types (KBM7 cells versus MiaPaCa-2 cells) and observed different mechanisms ^1, 2^. Besides PDEδ, several independent studies carried out by different groups on different targets including NAMPT, BRD4, CDK9, PSCK9, alpha-synuclein, lipid droplets, and mitochondria using GW5074 or other chemicals that we reported as LC3B binders, all suggested autophagy-mediated degradation mechanisms ^3, 4, 5, 6, 7, 16, 17, 18, 19^, confirming the feasibility of ATTEC, at least conceptually. Noticeably, several studies have shown that LD•ATTECs degrade lipid droplets ^7, 20, 21^, which are unlikely to be degraded by the DCAF11-proteasome pathway. It is possible that the targets that we selected to test in our previous studies, including the mHTT protein with expanded polyQ, the lipid droplets, and the mitochondria, are all difficult to be degraded by the proteasome and are capable of binding to multiple compound molecules (1:N stoichiometry). Thus, these targets may favor autophagic degradation mechanisms engaged by these compounds. On the other hand, in light of the data presented in ref 1, alternative mechanisms mediating the degradation via DCAF11 for certain targets in some cell types are also likely.

Taken together, we speculate that GW5074, AN1, and/or similar arylidene-indolinones may function as chemical “warheads” that may engage different degradation machinery when degrading different targets in different cell types. On one hand, this may give flexibility allowing a wider spectrum of degradation targets compared to traditional degrader technologies. On the other hand, stringent mechanistic experiments are required to elucidate the degradation mechanisms case-by-case for degraders with this type of warheads.

## Methods

### Surface Plasmon Resonance

SPR was performed using BIAcore 8K (Cytiva) at 25 °C. LC3B protein was diluted in 10 mM NaAc (pH 4.5, Cytiva, cat. no. BR100350) and immobilized onto CM5 sensor chip (Cytiva, cat. no. BR100530) using the standard amine coupling procedure (Cytiva, cat. no. BR100050) with a ~7000 units (RU) immobilization level. Compounds were diluted in PBS-P buffer (Cytiva, cat. no. 28995084) with a final concentration of 5% DMSO. Solvent correction was performed using a linear gradient of DMSO solution (4.50%, 5.05%, 5.24% and 5.8% DMSO in 1X PBS-P) to eliminate the influence of DMSO in the system. Affinity determination experiments were performed using a built-in multi-cycle mode. Data analysis was performed after solvent correction using the BIAcore 8K Insight Evaluation software (Cytiva). The affinity values (K_d_) were obtained by the steady-state affinity method.

### Biolayer Interferometry assay (BLI)

BLI assays were performed using ForteBio Octet R8 instrument (ForteBio, Inc., Menlo Park, USA). Purified LC3B was biotinylated using biotinylation kit (Genemore, Lot#1828M) and loaded onto super streptavidin biosensors (ForteBIO, cat. no. 18-5065-5057). Association-dissociation cycles of compounds were performed by moving and dipping LC3 loaded and reference sensors into compound solution wells (double diluted GW5074/AN1, 20 mM HEPES pH 7.4, 150 mM NaCl, 0.02% Tween 20, 2% DMSO) followed by moving into running buffer wells (20 mM HEPES pH 7.4, 150 mM NaCl, 0.02% Tween 20, 2% DMSO). The signals were then analyzed using a double reference subtraction protocol to eliminate nonspecific background signals and signal drifts caused by biosensor variability, and then fitted with a 1:1 stoichiometric model. A global fit of the steady-state signals was performed using the manufacturer’s protocol to calculate the K_d_.

### NMR spectroscopy

2D HSQC of titration experiments were collected on an Agilent 800 MHz spectrometer equipped with a cryogenic probe. All NMR experiments were collected at 298 K. 50 μM ^15^N-labelled LC3B protein was titrated by 500 μM molecule GW5074 and AN1 respectively in the NMR buffer of 50 mM Na_2_HPO_4_, 50 mM NaCl, 10% (vol/vol) DMSO and 10% (vol/vol) D_2_O at pH 7.0. All NMR data were analyzed with software Sparky.

### The Cellular Thermal Shift Assay (CETSA)

HeLa cells were seeded in 10 cm culture dishes with a 5×10^6^ cells/well density. 24 hours (h) later, cell pellets were collected and lysed at 4 °C for 1 h in 1x PBS+1% Tritonx-100+1x Complete protease inhibitor (Sigma-Aldrich, cat. no. 11697498001), and centrifuged at > 20,000x g at 4 °C for 10 min. Compounds (10 μM AN1 or 50 μM GW5074) or vehicle control (DMSO) were incubated with HeLa lysate supernatant for 2 h at room temperature. The respective lysates were divided into smaller (60 μL) aliquots and heated individually at different temperatures for 5 min with an AB Applied Biosystems Veriti 96 well cycler followed by cooling for 3 min at room temperature. The heated lysates were centrifuged at 20,000x g for 20 min at 4 °C to separate the soluble fractions from precipitates. The supernatants were transferred to new microtubes and analyzed by SDS-PAGE followed by Western blot analysis using anti-LC3B antibody (Thermo Fisher Scientific, cat. no. PA1-16930).

### Compound treatment in cells

The compounds used in this study were all commercially available, and quality controlled by the vendors using LS-MASS and NMR. GW5074: 3-3-((3,5-dibromo-4-hydroxyphenyl)methylidene)-5-iodo-1H-indol-2-one (MCE; cat. no. HY-10542); AN1: 5-bromo-3-[(4-hydroxyphenyl)methylidene]-2,3-dihydro-1H-indol-2-one (MCE; cat. no. HY-130258); MG-132 (Selleck, cat. no. S2619); CQ: chloroquine (Selleck, cat. no. S6999). Before compound treatment, HEK293T cells were plated at 8×105/well in a 6-well plate. After cell adhesion, cells were transfected with mHTTexon1-Q72-GFP plasmid and expressed for 24 h before compound treatment. For compound treatment in the cells, the compounds were first dissolved in DMSO to a 10-50 mM stock solution depending on the solubility. The stock solution was then serially diluted in DMSO and then to the culture medium to reach final concentration (typically 0.1~0.2% final DMSO in the culture). For CQ or MG-132 treated group, CQ or MG-132 were pretreated for 4 h before GW5074 or AN1 treatment to ensure the inhibition of autophagy or proteasome.

### Protein extraction and Western Blot

Cell pellets were collected and lysed at 4 °C for 1 h in 1x PBS+1% Tritonx-100+1x Complete protease inhibitor (Sigma-Aldrich, cat. no. 11697498001). The lysates were then centrifuged at 12,000x g at 4 °C for 15 min. The supernatants’ concentrations were quantified using the BCA method (Beyotime, cat. no. P0012). The remaining supernatants were transferred into loading samples: 65% supernatants; 10% DTT; 25% NuPAGE™ LDS Sample Buffer (4X) (Thermo Fisher Scientific, cat. no. NP0008) and were heated at 75 °C for 10 min. The loading samples were loaded in SDS-PAGE and transferred onto nitrocellulose membranes for Western blots.

### Real-time measurements of LC3B binding kinetics with mHTT-Q72 in the presence or absence of compounds

To measure the binding kinetics between mHTT-Q72 and LC3B in the presence or absence of GW5074 or AN1, recombinant purified LC3B was immobilized onto epoxy functionalized glass slides at a concentration of 35.7 μM. Each glass slide contains 24 identical protein microarrays, with each protein printed 10 times on a single microarray. The printed protein microarray was then assembled into a fluidic cartridge and the slide was washed in situ with HEPES to remove any excess unbound samples. After washing, the slide was blocked with 7,600 nM BSA in HEPES for 10 min.

For the binary binding kinetics measurement, three different concentrations of recombinant purified mHTT-Q72 (200 nM, 800 nM, 1600 nM) were flowed over three separate fresh microarrays. In the ternary binding experiment, different concentrations of mHTT-Q72 (200 nM, 400 nM, 800 nM, 1600 nM) were preincubated with GW5074 or AN1 at concentrations two times that of mHTT-Q72 or forty times that of mHTT-Q72 for 2 h. After preincubation, the mixtures were flown over three separate fresh protein microarrays.

Both the binary and ternary binding kinetics were recorded using a scanning OI-RD microscope. The reaction kinetic rate constants were extracted by fitting the binding curves globally using the one-to-one Langmuir reaction model.

